# Purifying selection can reduce intraspecific mitochondrial gene variation to that of nuclear rRNA

**DOI:** 10.1101/2020.03.31.017764

**Authors:** Tshifhiwa G. Matumba, Jody Oliver, Nigel P. Barker, Christopher D. McQuaid, Peter R. Teske

## Abstract

Mitochondrial DNA (mtDNA) has long been used to date the divergence between species, and to explore the time when species’ effective population sizes changed. The idea that mitochondrial DNA is useful for molecular dating rests on the premise that its evolution is neutral. This premise was questionable to begin with, and even though it has long been challenged, the evidence against clock-like evolution of mtDNA is usually ignored. Here, we present a particularly clear and simple example to illustrate the implications of violations of the assumption of selective neutrality. DNA sequences were generated for the mtDNA COI gene and the nuclear 28S rRNA of two closely related and widely distributed rocky shore snails whose geographical ranges are defined by different thermal preferences. To our knowledge, this is the first study to use nuclear rRNA sequence for studying species-level genealogies instead of phylogenetics, presumably because this marker is considered to be uninformative at this taxonomic level. Even though the COI gene evolves at least an order of magnitude faster, which was reflected in high inter-specific divergence, intraspecific genetic variation was similar for both markers. As a result, estimates of population expansion times based on mismatch distributions were completely different for the two markers. Assuming that 28S evolves effectively clock-like, these findings likely illustrate variation-reducing purifying selection in mtDNA at the species level, and an elevated divergence rate caused by divergent selection between the two species. Although these two selective forces together make mtDNA suitable as a DNA barcoding marker because they create a ‘barcoding gap’, estimates of demographic change can be expected to be highly unreliable. Our study contributes to the growing evidence that the utility of mtDNA beyond DNA barcoding is limited.

## Introduction

Mitochondrial DNA (mtDNA) has long been a marker of choice for investigating concepts as diverse as estimating genetic diversity and effective population sizes, reconstructing species’ evolutionary histories, exploring spatial genetic subdivisions, and identifying cryptic species. All these methods assume that mtDNA variation conforms to the neutral model of molecular evolution (Kimura 1983), but violations of this premise have long been recognised (Ballard and Kreitman 1995). Over the past decades, much evidence has accumulated that mtDNA can be strongly affected by selective sweeps and background selection (Blier et al. 2001; Ballard and Rand 2005; Meiklejohn et al. 2007; Stewart et al. 2008). As a result, the usefulness of the marker in assessing genetic diversity (Bazin et al. 2006) and exploring spatial genetic structure in continuously distributed populations (Teske et al. 2018) has been questioned, and corrections of the mitochondrial molecular clock that account for selection have been proposed (Soares et al. 2009).

The implications of reduced genetic diversity at the species or population levels due to purifying selection has so far received little attention. When mutations in mitochondrial genes occur at fewer sites than expected under the neutral model (Lawrie et al. 2013), molecular dating of historical demographic events by means of evolutionary rate estimates that are typically based on inter-specific divergence (Knowlton and Weigt 1998; Schubart et al. 1998) will result in considerable underestimates. This is particularly likely because divergence between species can be strongly affected by diversifying selection that is driven by different environmental conditions (Lamb et al. 2018; Sun et al. 2018), resulting in a faster accumulation of mutations characterising each species than is expected under the neutral model.

Here, we explore this issue using mitochondrial and nuclear DNA sequence data from two common southern African snails of the genus *Afrolittorina*. The finding that data from two gene regions whose mutation rates are assumed to differ by at least an order of magnitude have similar levels of intraspecific variation challenges the usefulness of mitochondrial DNA sequences for studying historical demographic changes.

## Materials and Methods

The two study species, *Afrolittorina africana* (Philippi, 1847) and *A. knysnaensis* (Philippi, 1847) are the most widespread and abundant littorinid snails on rocky shores along the southern African coast (McQuaid 1992). They were formerly believed to comprise a single species whose morphological appearance gradually changes along its range (Hughes 1979), but their species status is now confirmed using both molecular and morphological data (Reid and Williams 2004).

Specimens of *Afrolittorina* were collected from 34 localities across the species’ distribution ranges along the southern African coastline (supplemental online material, Table S1). Neither species exhibits genetic structure throughout its range (Matumba 2013). Laboratory procedures are described in the supplemental online material, and the primers used for PCR amplification of the mtDNACOI gene and the nuclear 28S rRNA are listed in Table S2.

Genealogical relationships between COI haplotypes and 28S alleles were reconstructed using the median joining algorithm (Bandelt et al. 1999) in popArt v1.7 (Leigh and Bryant 2015). Sites containing gaps were excluded. Variable sites in the mtDNA sequences were explored for evidence of intra- and inter-specific selection using a McDonald-Kreitman test (McDonald and Kreitman 1991) in MKT (Egea et al. 2008). The invertebrate mitochondrial code was specified, and the Jukes-Cantor model (Jukes and Cantor 1969) was used to correct for divergence. To explore the effect of using interspecific evolutionary rates to estimate when intraspecific demographic changes occurred, we calculated population expansion time under the sudden-expansion model (Rogers and Harpending 1992) using Arlequin v3.5 (Excoffier and Lischer 2010). For each marker type, we selected the slowest and the fastest published marine gastropod rates from the literature. For COI, these were 0.5%.my^-1^ (Malaquias and Reid 2009) and 2.6%.my^-1^ (Williams and Reid 2004), respectively. For 28S, a slow rate of 0.01%.my^-1^ (Malaquias and Reid 2009) and a fast rate of 0.05%.my^-1^ (Williams and Reid 2004) were used.

## Results

Species-specific genetic clusters were highly distinct (Fig. 1a), with a minimum number of 44 nucleotide differences between the two species’ most closely related haplotypes. In contrast to the COI sequences, differentiation between 28S sequences (Fig. 1b) was an order of magnitude smaller (4 nucleotide differences).

**Fig. 1.**
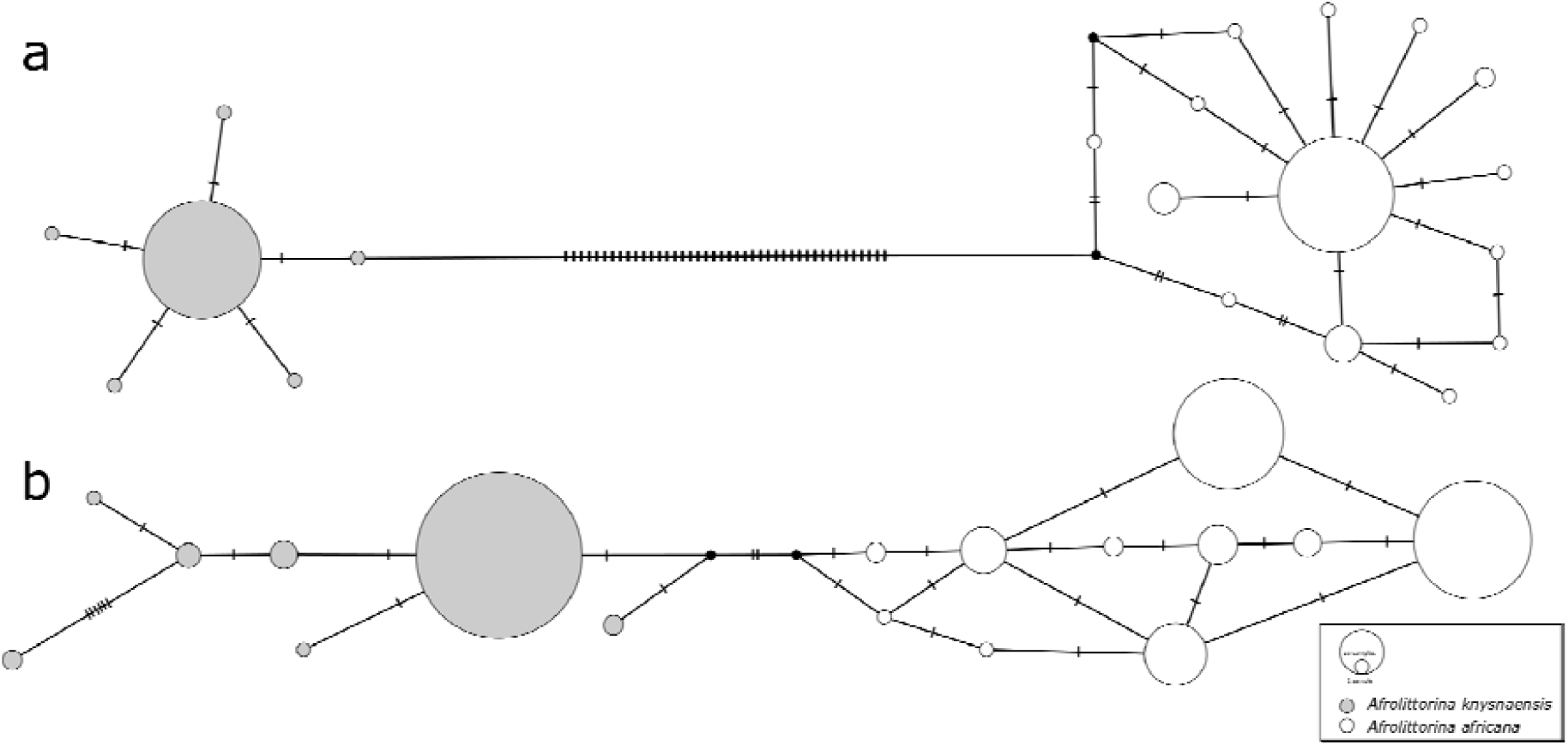
Median-joining haplotype networks constructed from a) COI sequences of b) 28S rRNA sequences of *Afrolittorina knysnaensis* (grey) and *A. africana* (white). Low intra-specific variation and high inter-specific variation of COI potentially illustrate stabilising and divergent selection, respectively. The size of circles is proportional to the frequency of each haplotype, cross-bars represent nucleotide differences, and black dots are missing haplotypes not found in the samples.

In contrast to the high inter-specific differentiation between COI haplotypes, intra-specific genetic differentiation was comparatively low for this marker, and similar to that of 28S. In *A. knysnaensis*, six COI haplotypes and seven 28S haplotypes were found, while the maximum differentiation between the COI haplotypes was only two nucleotide differences, but 10 for 28S. The number of haplotypes was greater for *A. africana*, where 14 were found for COI and 10 for 28S. Maximum nucleotide differences for this species were seven in the COI network and five for 28S.

All five variable sites in *A. knysnaensis*, and 11 out of 12 variable sites in *A. africana*, were found at synonymous third character positions. The remaining site in *A. africana* was at a first character position but was also synonymous (CUG vs. UUG, which both code for Leu using the invertebrate mitochondrial code). For the complete dataset, which comprised 58 mutations of which the majority was found between species, 57 mutations were synonymous and one was non-synonymous. The McDonald-Kreitman test had the highest possible proportion of adaptive substitutions (α) of 1.0, but was nonetheless non-significant (χ^2^ = 0.33, P = 0.57), supposedly because of the lack of non-synonymous polymorphism.

The practical implications of two markers with very different evolutionary rates based on inter-specific divergence having similar levels of intraspecific variation are illustrated in Table 1. Using published rates, estimates of population expansion times were several orders of magnitude greater for the 28S data than for the COI data.

**Table 1.** Estimates of population expansions of the two species of *Afrolittorina* under the sudden expansion model. The moment estimator τ is equal to 2ut, where u equals 2 µk (μ is the mutation rate and k is the length of the sequence), and *t* is the time of expansion.

| Species | $\tau$ | Marker | $\mu$ (%.my<sup>-1</sup>) | $t$ (my) |
| --- | --- | --- | --- | --- |
| *Afrolittorina knysnaensis* | 2.00 | COI | 0.50<sup>1</sup> | 0.40 (0.00 – 0.41) |
|  |  |  | 2.60<sup>2</sup> | 0.07 (0.00 – 0.08) |
|  | 3.25 | 28S | 0.01<sup>1</sup> | 32.1 (18.5 – 61.3) |
|  |  |  | 0.05<sup>2</sup> | 6.41 (3.69 – 12.3) |
| *Afrolittorina africana* | 2.50 | COI | 0.50<sup>1</sup> | 0.50 (0.30 – 0.79) |
|  |  |  | 2.60<sup>2</sup> | 0.10 (0.06 – 0.15) |
|  | 2.75 | 28S | 0.01<sup>1</sup> | 27.1 (19.1 – 51.5) |
|  |  |  | 0.05<sup>2</sup> | 5.42 (3.81 – 10.3) |

<sup>1</sup>Malaquias & Reid 2009; <sup>2</sup>Williams & Reid 2004. A generation time of one year was used.

## Discussion

The usefulness of the mtDNA COI gene to uncover overlooked biodiversity is undisputed because of the marker’s tendency to have a well-defined barcoding gap (as was also found here). Many researchers explore their mtDNA data for additional information, but variation-reducing selection at the population or species levels, and adaptive selection at the inter-species level that together create the barcoding gap (Stoeckle and Thaler 2014) make the usefulness of mtDNA for other applications questionable (Bazin et al. 2006; Teske et al. 2018). The finding that only synonymous mutations were found at the intra-specific level in *Afrolittorina*, whereas inter-specific mutations included a non-synonymous mutation, is by no means strong evidence for intra-specific neutrality and inter-specific selection. Recent evidence shows that selection also affects synonymous mutations, and evidence for purifying selection at the species level typically comes from a smaller number of synonymous mutations than expected under the neutral model, rather than the presence of non-synonymous mutations (Lawrie et al. 2013). Of the different gene regions that make up the mitochondrial genome, COI and COII are under particularly strong purifying selection (Stewart et al. 2008). In the present study, we have highlighted a largely unexplored problem resulting from selection in mtDNA data: the fact that the reconstruction of demographic events that are potentially affected by purifying selection are typically dated using molecular clocks that were calibrated using variation-increasing inter-specific divergence.

The differences in variation found between the mtDNA COI sequences and the nrDNA 28S rRNA sequences of two closely related snails of the genus *Afrolittorina* likely reflect differences in the extent of selection pressure. The snails’ COI sequences were much more strongly differentiated than their 28S sequences, potentially reflecting diversifying selection as a result of adaption to different thermal environments. In contrast, there was comparatively little genetic variation at the intraspecific level for both markers. To our knowledge, this is the first study to use 28S to explore genetic variation at the species level, perhaps because it is commonly assumed that this marker evolves so slowly that it would only be informative at deeper taxonomic levels. The finding that the barcoding gap is not strongly developed in 28S rRNA suggests that the evolution of this marker is more likely to conform to the expectations of the neutral model of evolution.

This finding has important implications for the interpretation of species-level mtDNA data. Both species of *Afrolittorina* in the haplotype network have a ‘star-phylogeny’, where a dominant haplotype has given rise to a large number of comparatively rare haplotypes (Castelloe and Templeton 1994). Star-phylogenies are often interpreted as signatures of a population bottleneck that resulted in a reduction of genetic diversity until a single haplotype was left, for instance during a previous glacial phase (Magoulas et al. 1996), followed by a rapid increase in population size. The finding that the corresponding but similarly variable 28S haplotype networks are less star-like (although the *A. knysnaensis* network is more star-like than that of *A. africana*) suggests that species-level purifying selection could be an alternative explanation for the patterns found in the mtDNA data. In this case, the apparent population bottleneck would instead be attributed to a selective sweep that has reduced genetic diversity, which is expected to be common in species with very large population sizes (Gillespie 2001), such as littorinid snails.

Even if the assumption that demographic expansions have taken place is correct, the dating of these may be strongly affected by selection. Estimation of demographic trends by means of mismatch distributions using only mtDNA sequence data continue to be widely used in the recent literature (Low et al. 2017; Diringer et al. 2019; Gao et al. 2019; Iván Pérez-Quiñonez et al. 2019) although not all have used the method for molecular dating. A similar method, Bayesian skyline plots (Drummond et al. 2005), is also frequently used with mtDNA data only, and findings tend to be similar (Grant 2015).

The finding that intraspecific mtDNA variation can be as low as that of slowly-evolving nuclear rRNA cautions against the continued use of mtDNA for exploring historical demographic changes. In our opinion, it is time to discontinue the use of fixed mtDNA rates based on divergence dating of closely related taxa, such as the closure of the Central American Seaway to date phylogenies of marine species (Knowlton and Weigt 1998; Schubart et al. 1998) or the 2% rule in birds (Shields and Wilson 1987). The very large datasets generated using next-generation sequencing have considerable potential to facilitate more accurate dating by identifying nuclear markers that conform to the assumptions of a molecular clock. Curiously, fixed rates based on mtDNA data are still being used to calibrate such datasets when no suitable fossil calibration points exist (Trucchi et al. 2014). A possible solution in such cases may involve the identification of a suite of neutral markers that can be used to assess divergence between the species used in the original molecular dating studies, and 28S rRNA may be a suitable candidate.

## Acknowledgements

This work is based upon research supported by the National Research Foundation of South Africa (Grant number 64801) and Rhodes University.

